# The rise and fall of SARS-CoV-2 variants and the emergence of competing Omicron lineages

**DOI:** 10.1101/2022.02.09.479842

**Authors:** Tanner Wiegand, Aidan McVey, Anna Nemudraia, Artem Nemudryi, Blake Wiedenheft

## Abstract

In late December of 2019, high throughput sequencing technologies enabled rapid identification of SARS-CoV-2 as the etiological agent of COVID-19, and global sequencing efforts are now a critical tool for monitoring the ongoing spread and evolution of this virus. Here, we analyze a subset (n=83,204) of all publicly available SARS-CoV-2 genomes (n=~5.6 million) that were randomly selected, but equally distributed over the course of the pandemic. We plot the emergence and extinction of new variants of concern (VOCs) over time and show how this corresponds to the ongoing accumulation of mutations in SARS-CoV-2 genomes and individual proteins. While the accumulation of mutations generally follows a linear regression, non-synonymous mutations are significantly greater in Omicron viruses than in previous variants–especially in the spike and nucleoproteins–and these differences are more pronounced in a recently identified sub-lineage (BA.2) of Omicron.

**Importance:** Omicron is the fifth SARS-CoV-2 variant to be designated a Variant of Concern (VOC) by the World Health Organization (WHO). Here we provide a retrospective analysis of SARS-CoV-2 variants and explain how the Omicron variant is distinct. Our work shows that the spike and nucleoproteins have accumulated the most mutations in Omicron variants, but that the accessory proteins of SARS-CoV-2 sequences are changing most rapidly relative to their size. Collectively, this “Observation” provides a concise overview of SARS-CoV-2 evolution, reveals mutational differences between two Omicron lineages, and highlights changes in the SARS-CoV-2 proteome that have been under reported.

## Main Text

All viruses, including SARS-CoV-2, accumulate mutations as they replicate and spread. Most of these changes have little or no impact on transmissibility, disease severity, or the effectiveness of current vaccines and diagnostics. However, selective pressures that act on random mutations efficiently enrich rare variants with enhanced viral fitness (e.g., replication and transmissibility) (1,2). Initially, SARS-CoV-2 variants were named according to their geographic origin (i.e., Wuhan), but naming based on geography can be culturally insensitive, stigmatizing, and is often inaccurate. Thus, this naming scheme was quickly replaced by a unique combination of letters and numbers known as the Pangolin nomenclature (e.g., B.1.1.7), but this appellative is sometimes confusing for scientists and the public alike (3). With increasing frequency, we now complement the Pangolin nomenclature with the use of a single Greek letter. Omicron (BA.1 or B.1.1.529.1) is the 15^th^ letter of the Greek alphabet, but only the fifth variant designated as a Variant of Concern (VOC) by the World Health Organization (WHO).

Omicron was first identified from a specimen collected on November 9^th^ of 2021 in South Africa, and was designated as a VOC on November 26^th^ (4). This designation is based on the number of mutations (26–32) in the spike protein relative to previously sequenced isolates, as well as concerning epidemiological reports from South Africa (5, 6). Importantly, real-time tracking of pathogen evolution by the team at Nextstrain (7), has recently identified an offshoot lineage of Omicron (i.e., BA.2 or B.1.1.529.2) that has twenty shared and six unique spike mutations compared to archetypal Omicron (i.e., BA.1) viruses. However, the spike is only one of at least 26 SARS-CoV-2 encoded proteins (8). Here we analyze non-synonymous and synonymous mutations in SARS-CoV-2 genomes and highlight how different proteins evolve at distinct rates (**Figure 1**).

**Figure 1.**
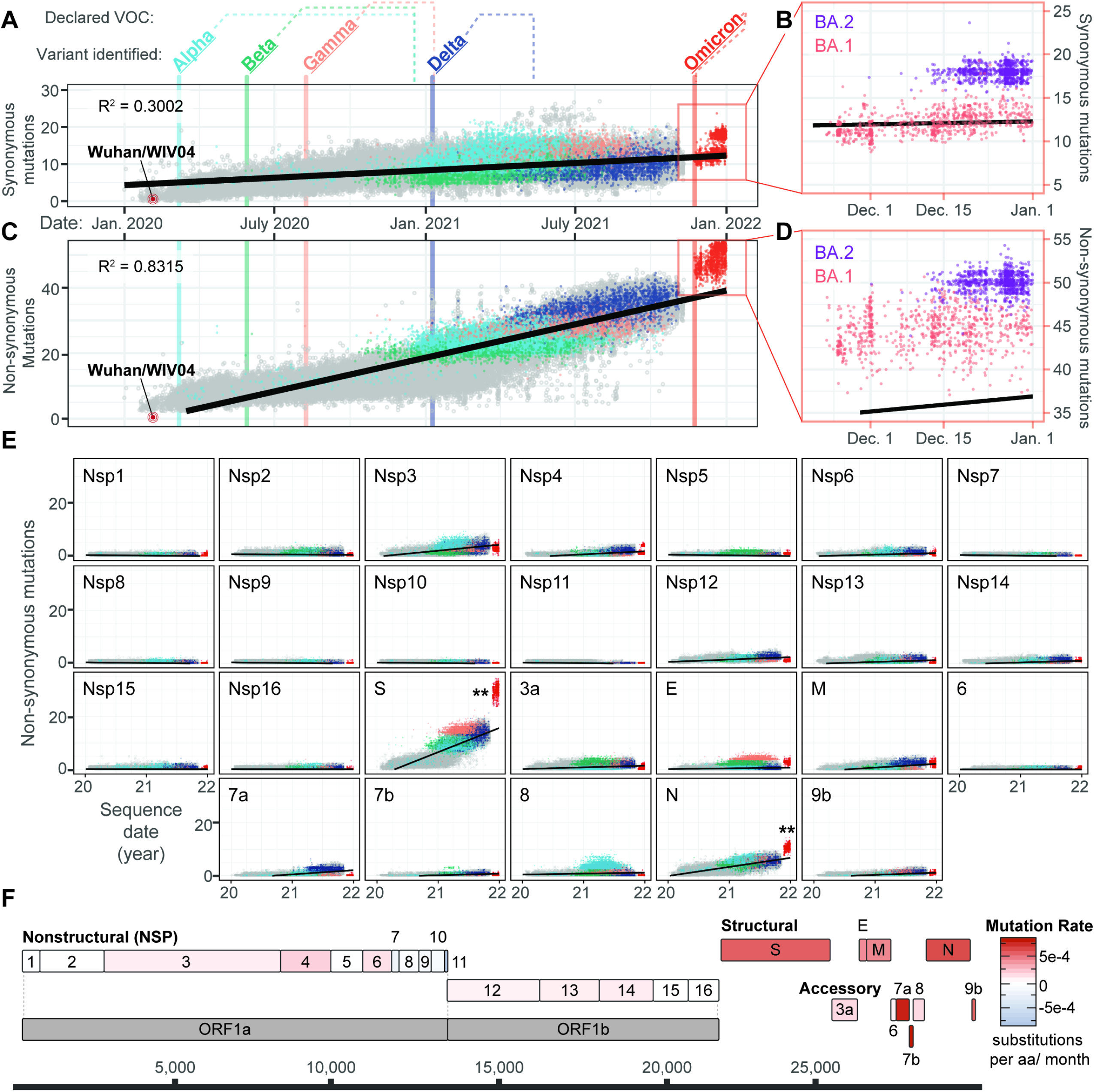
Evolution of SARS-CoV-2 and the emergence of new variants. **A)** Synonymous mutations in 83,204 genomes sampled from Dec 19, 2021 to Jan 14, 2022 (GISAID accessions available at: github.com/WiedenheftLab/Omicron). Variants of concern (VOC) are labeled in bold, with dates of the first sequence for each lineage shown as vertical lines through the graph. The time elapsed between first detection and VOC designation by the WHO is shown as dotted lines above the graph. Dots are colored similar to variant names, and grey circles represent non-VOC lineage genomes. A linear regression line is shown in black. Omicron genomes are <2.58 sigma from the mean, indicating that do not significantly deviate from the expected number of mutations. **B)** Inset from Panel A. Omicron is divided into two lineages, BA.1(red) and BA.2 (purple). Synonymous mutations are significantly different between these two lineages (T-test, p < 2e-16). **C)** Non-synonymous mutations in SARS-CoV-2 genomes are plotted as in panel A. Omicron variants deviate from the linear trendline (i.e., expected mutation rate) as the majority are >2.58 sigma from the mean (p < 0.01). **D)** Inset from Panel C. Non-synonymous in BA.1 (red) are significantly different from BA.2 (purple) (T-test, p < 2e-16). **E)** Non-synonymous mutations accumulated in SARS-CoV-2 proteins over time. Linear regression lines and variant colors are plotted as in panel A. Double-asterisks denote statistically significant (p < 0.01) increases in Omicron mutation rates for a given protein, compared to mutation counts predicted by linear regression analysis of non-Omicron sequences. **F)** Schematic depiction of SARS-CoV-2 protein coding sequences, with each gene colored according to non-synonymous rates predicted in panel D that have been normalized to the length of each respective protein.

First, we filtered all SARS-CoV-2 genomes available on GISAID (n= ~5.6 million on December 2, 2021) for a subset of genomes that had fewer than 3,000 ambiguous base calls and were considered high-coverage by GISAID (9). The remaining genomes were further pruned using NextClade (7). Only sequences with good quality scores were retained, which resulted in ~1.7 million “high-quality” genomes. We then compared randomly selected “high-quality” genomes that were equally distributed over the course of the pandemic; starting with the Wuhan reference genome and ending with all high-quality Omicron genomes available on GISAID (n=2,136, on January 14, 2022) (9). This analysis reveals a steady increase in synonymous mutations over time (**Figure 1A**). While the number of synonymous mutations in BA.1 and BA.2 lineages of Omicron are not significantly different from what would be expected based on a simple linear regression, BA.2 has significantly more mutations than BA.1 (**Figure 1B)**. A similar, but more pronounced pattern is also evident for the number of non-synonymous mutations, which are significantly elevated in Omicron genomes (**Figure 1C and D**).

To understand which gene or genes, contribute most to the significant jump in non-synonymous mutations in Omicron, we analyzed each of the SARS-CoV-2 genes individually. This analysis reveals two genes that have significantly more non-synonymous mutations in both BA.1 and BA.2 viruses. The spike protein of Omicron has the most changes (24-35 mutations, mean = 30.7) (**Figure 1E**), and the impact of these mutations on viral transmission, vaccination and effectiveness of diagnostics has been widely discussed (10–12). However, the nucleoprotein (N) of Omicron viruses also has significantly more non-synonymous mutations, which suggests that diagnostics targeting this protein may also become less efficacious in the future. Indeed, the American Food and Drug Administration (FDA) has already warned of possible decreases in the sensitivity of some COVID-19 tests that rely on detection of the nucleoprotein, due to mutations in Omicron variants (13).

While S and N proteins account for the most mutations, this analysis does not account for differences in protein length. To account for length, we normalized non-synonymous mutation rates determined in **Figure 1E** by dividing each by the number of amino acids present in their respective proteins (**Figure 1F**). While changes in the spike attract the most public attention, accounting for protein length reveals that SARS-CoV-2 accessory proteins (i.e., 7a, 7b, and 9b) are changing most rapidly. These viral accessory proteins are implicated in innate immune suppression (14, 15) and recent evidence suggest that mutations in these proteins may play important (and somewhat underappreciated) roles in viral evolution (16, 17).

## Acknowledgments

Research in the Wiedenheft lab is supported by the National Institutes of Health (1R35GM134867), the Montana State University (MSU) Agricultural Experimental Station, the MJ Murdock Charitable Trust, the Gianforte Foundation, the State of Montana, the City of Bozeman, a charitable donation from the Rosolowsky family, and the MSU Office of the Vice President for Research. We thank the GISAID’s EpiFlu Database and contributing laboratories. The analyses in this paper would not have been possible without their willingness to share data. Accessions of sequence utilized in this work are accessible at github.com/WiedenheftLab/Omicron.

## References

1. Ramesh S, Govindarajulu M, Parise RS, Neel L, Shankar T, Patel S, Lowery P, Smith F, Dhanasekaran M, Moore T. 2021. Emerging SARS-CoV-2 Variants: A Review of Its Mutations, Its Implications and Vaccine Efficacy. Vaccines (Basel) 9.

2. Redd AD, Nardin A, Kared H, Bloch EM, Pekosz A, Laeyendecker O, Abel B, Fehlings M, Quinn TC, Tobian AAR. 2021. CD8+ T-Cell Responses in COVID-19 Convalescent Individuals Target Conserved Epitopes From Multiple Prominent SARS-CoV-2 Circulating Variants. Open Forum Infect Dis 8:ofab143.

3. Rambaut A, Holmes EC, O’Toole A, Hill V, McCrone JT, Ruis C, du Plessis L, Pybus OG. 2020. A dynamic nomenclature proposal for SARS-CoV-2 lineages to assist genomic epidemiology. Nat Microbiol 5:1403–1407.

4. WHO. 2021. Classification of Omicron (B.1.1.529): SARS-CoV-2 Variant of Concern. https://www.whoint/news/item/26-11-2021-classification-of-omicron-(b11529)-sars-cov-2-variant-of-concern.

5. Chen J, Wang R, Gilby NB, Wei GW. 2021. Omicron (B.1.1.529): Infectivity, vaccine breakthrough, and antibody resistance. ArXiv.

6. Callaway E, Ledford H. 2021. How bad is Omicron? What scientists know so far. Nature 600:197–199.

7. Hadfield J, Megill C, Bell SM, Huddleston J, Potter B, Callender C, Sagulenko P, Bedford T, Neher RA. 2018. Nextstrain: real-time tracking of pathogen evolution. Bioinformatics 34:4121–4123.

8. Finkel Y, Mizrahi O, Nachshon A, Weingarten-Gabbay S, Morgenstern D, Yahalom-Ronen Y, Tamir H, Achdout H, Stein D, Israeli O, Beth-Din A, Melamed S, Weiss S, Israely T, Paran N, Schwartz M, Stern-Ginossar N. 2021. The coding capacity of SARS-CoV-2. Nature 589:125–130.

9. Elbe S, Buckland-Merrett G. 2017. Data, disease and diplomacy: GISAID’s innovative contribution to global health. Glob Chall 1:33–46.

10. Saxena SK, Kumar S, Ansari S, Paweska JT, Maurya VK, Tripathi AK, Abdel-Moneim AS. 2021. Characterization of the novel SARS-CoV-2 Omicron (B.1.1.529) variant of concern and its global perspective. J Med Virol doi:10.1002/jmv.27524.

11. Lupala CS, Ye Y, Chen H, Su XD, Liu H. 2022. Mutations on RBD of SARS-CoV-2 Omicron variant result in stronger binding to human ACE2 receptor. Biochem Biophys Res Commun 590:34–41.

12. Harvey WT, Carabelli AM, Jackson B, Gupta RK, Thomson EC, Harrison EM, Ludden C, Reeve R, Rambaut A, Consortium C-GU, Peacock SJ, Robertson DL. 2021. SARS-CoV-2 variants, spike mutations and immune escape. Nat Rev Microbiol 19:409–424.

13. FDA. 2021. SARS-CoV-2 Viral Mutations: Impact on COVID-19 Tests. Omicron Variant: Impact on Antigen Diagnostic Tests (As of 12/28/2021). https://www.fdagov/medical-devices/coronavirus-covid-19-and-medical-devices/sars-cov-2-viral-mutations-impact-covid-19-tests#omicronvariantimpact.

14. Thorne LG, Bouhaddou M, Reuschl AK, Zuliani-Alvarez L, Polacco B, Pelin A, Batra J, Whelan MVX, Hosmillo M, Fossati A, Ragazzini R, Jungreis I, Ummadi M, Rojc A, Turner J, Bischof ML, Obernier K, Braberg H, Soucheray M, Richards A, Chen KH, Harjai B, Memon D, Hiatt J, Rosales R, McGovern BL, Jahun A, Fabius JM, White K, Goodfellow IG, Takeuchi Y, Bonfanti P, Shokat K, Jura N, Verba K, Noursadeghi M, Beltrao P, Kellis M, Swaney DL, Garcia-Sastre A, Jolly C, Towers GJ, Krogan NJ. 2021. Evolution of enhanced innate immune evasion by SARS-CoV-2. Nature doi:10.1038/s41586-021-04352-y.

15. Young BE, Fong SW, Chan YH, Mak TM, Ang LW, Anderson DE, Lee CY, Amrun SN, Lee B, Goh YS, Su YCF, Wei WE, Kalimuddin S, Chai LYA, Pada S, Tan SY, Sun L, Parthasarathy P, Chen YYC, Barkham T, Lin RTP, Maurer-Stroh S, Leo YS, Wang LF, Renia L, Lee VJ, Smith GJD, Lye DC, Ng LFP. 2020. Effects of a major deletion in the SARS-CoV-2 genome on the severity of infection and the inflammatory response: an observational cohort study. Lancet 396:603–611.

16. Gordon DE, Jang GM, Bouhaddou M, Xu J, Obernier K, White KM, O’Meara MJ, Rezelj VV, Guo JZ, Swaney DL, Tummino TA, Huttenhain R, Kaake RM, Richards AL, Tutuncuoglu B, Foussard H, Batra J, Haas K, Modak M, Kim M, Haas P, Polacco BJ, Braberg H, Fabius JM, Eckhardt M, Soucheray M, Bennett MJ, Cakir M, McGregor MJ, Li Q, Meyer B, Roesch F, Vallet T, Mac Kain A, Miorin L, Moreno E, Naing ZZC, Zhou Y, Peng S, Shi Y, Zhang Z, Shen W, Kirby IT, Melnyk JE, Chorba JS, Lou K, Dai SA, Barrio-Hernandez I, Memon D, Hernandez-Armenta C, et al. 2020. A SARS-CoV-2 protein interaction map reveals targets for drug repurposing. Nature 583:459–468.

17. Nemudryi A, Nemudraia A, Wiegand T, Nichols J, Snyder DT, Hedges JF, Cicha C, Lee H, Vanderwood KK, Bimczok D, Jutila MA, Wiedenheft B. 2021. SARS-CoV-2 genomic surveillance identifies naturally occurring truncation of ORF7a that limits immune suppression. Cell Rep 35:109197.

